# Thalamic input drives co-timed excitation and inhibition to suppress cortical neuronal variability during movement initiation

**DOI:** 10.64898/2026.05.21.722630

**Authors:** Julian Ammer, Joshua Dacre, Julia Schiemann, Sara Moberg, Nathalie Rochefort, Matthias Hennig, Ian Duguid

## Abstract

Cortical activity is inherently dynamic, with ongoing fluctuations introducing variability into sensory-to-motor transformations. During movement initiation, neural variability in the motor cortex is sharply reduced, or “quenched,” a general phenomenon thought to ensure reliable population responses across cortical areas. However, the synaptic and circuit-level basis for this reduction in neural variability remains unclear. Here, we show that excitatory and inhibitory neurons in layer 5B of mouse motor cortex are co-activated during movement initiation in a cued forelimb push task. Co-timed excitation and inhibition drive pyramidal neuron membrane potential (*V_m_*) trajectories towards the compound synaptic reversal potential, stabilizing the *V_m_* and reducing trial-by-trial variability. Gain- and loss-of-function experiments demonstrate that thalamocortical input is critical for the coordinated recruitment of both excitation and inhibition and variability quenching at the cellular level, while also driving a cortical network state permissive for the expression of reproducible output dynamics. Our findings uncover a simple yet fundamental feedforward mechanism where thalamic input drives neural variability quenching to ensure reliable, structured population dynamics during movement initiation.

## Introduction

Motor cortical activity exhibits trial-by-trial variability during movement execution, a feature displayed by single neurons and neural populations (Churchland et al., 2006; Churchland & Shenoy, 2007; Fortier et al., 1993; Lee et al., 1998; Riehle et al., 2018; Riehle & Requin, 1993; Saberi-Moghadam et al., 2016). Far from being noise, this variability is crucial for behavioral flexibility and learning (Dhawale et al., 2017; Mandelblat-Cerf et al., 2009; Renart & Machens, 2014; Waschke et al., 2021). However, during movement initiation, a profound suppression – or quenching – of neuronal variability emerges, which is thought to ensure reliable cortical output during the execution of learned, skilled movements (Churchland et al., 2006, 2010; Kondapavulur et al., 2022; Makino et al., 2017; Rickert et al., 2009; Riehle et al., 2018). Despite the across species prevalence of this phenomenon, the biophysical mechanisms underpinning variability quenching in motor cortex remain unknown. While there is a paucity of empirical insights, theoretical work has proposed diverse network mechanisms capable of quenching variability in both sensory and motor regions (Bujan et al., 2015; Deco & Hugues, 2012; Guo & Kumar, 2023; Hennequin et al., 2018; Litwin-Kumar & Doiron, 2012; Rostami et al., 2024). These models converge on a key prediction: disrupting long-range input to that region should attenuate variability quenching (Goris et al., 2024). The idea being that this input drives both activity in individual target neurons and, through recurrent connectivity, triggers co-activation of local excitatory and inhibitory populations to drive variability quenching in the local network, rather than being purely inherited from upstream (Goris et al., 2024; Guo & Kumar, 2023; Hennequin et al., 2018; Rostami et al., 2024; Rubin et al., 2015). What remains unclear is what drives coordinated E-I activation and how this leads to neuronal variability quenching. Two possible sources of long-range input are interareal corticocortical projections or feedforward input from thalamus (Alyahyay et al., 2023; Goz & Hooks, 2023; Nashef et al., 2022; Okoro et al., 2022). Since both bottom-up and top-down intracortical inputs increase, rather than decrease, variability (Gómez-Laberge et al., 2016) long-range thalamocortical input remains the most likely candidate.

Thalamocortical input to primary motor cortex is essential for both the initiation and coordination of limb movements in rodents generating membrane potential (*V_m_*) changes that bidirectionally modulate layer 5 output (Dacre et al., 2021; Sauerbrei et al., 2020; Tanaka et al., 2018). Since thalamic afferents target both excitatory and inhibitory neurons, driving balanced network dynamics via intralaminar and translaminar crosstalk (Anderson et al., 2010; Apicella et al., 2012; Dura-Bernal et al., 2023; Hooks et al., 2013; Nashef et al., 2022; Tanaka et al., 2011; Weiler et al., 2008), it remains the most likely candidate to drive variability quenching during movement initiation (Churchland et al., 2006, 2010; Kondapavulur et al., 2022; Makino et al., 2017; Rickert et al., 2009; Riehle et al., 2018).

To test this idea, we performed intracellular and high-density extracellular probe recordings in mice trained to perform a cued forelimb lever push task. During movement initiation we observed co-activation of inhibitory and excitatory populations in layer 5B of motor cortex and a concomitant quenching of cortical output variability. This co-activation was reflected in the synaptic inputs to individual layer 5B pyramidal neurons, where concurrent excitation and inhibition generated a profound increase in the compound synaptic conductance. By combining our neuronal recordings with conductance-based modeling, we demonstrated that the increase in synaptic conductance consistently drives the *V_m_* towards the synaptic reversal potential, independent of the trial-by-trial baseline variability. Crucially, we identified thalamocortical input as a critical driver of the conductance increase, necessary for quenching single neuron *V_m_* variance and generating consistent across-trial population-based output during movement initiation.

## Results

### Coincident activation of excitatory and inhibitory neurons correlates with neuronal variability quenching

To investigate the mechanism underlying variability quenching, we trained mice to push a horizontal lever (4 mm) in response to a 6-kHz auditory cue for reward (Dacre et al., 2021). Spontaneous pushes during the inter-trial interval (4–6 s, randomized), partial pushes, or misses resulted in a lever reset to the start position without reward (Figures 1A). Mice rapidly learned the task achieving high success rates (proportion of successful pushes = 0.95, 95% CI [0.76 0.98], N = 9 mice, Figure 1B) with moderate reaction times (median RT = 270 ms 95% CI [197 563] ms, median IQR = 97 ms 95% CI [40 236] ms , N = 9 mice, ,Figure 1C). We investigated neuronal variability quenching during task execution by analyzing previously acquired high-density extracellular recordings from putative layer 5B pyramidal neurons in CFA of motor cortex (spike width > 0.4 ms, n = 323 neurons from N = 9 mice, Figures 1D-1E) (Currie et al., 2022; Dacre et al., 2021). To quantify firing rate variability, we calculated the population Fano factor (i.e. linear fit to the spike-count variance vs the spike-count mean of all layer 5B pyramidal neurons in a 100ms sliding window) (Churchland et al., 2010). Variability quenching (apparent as a reduction in Fano factor) began approximately 200 ms before and peaked at the point of movement initiation (Figure 1F). This effect was not due to the higher firing rates around movement onset as quenching was similar for mean matched data in which data points are randomly omitted to match the distribution of firing rates across time (see Methods and Churchland et al., 2010).

**Figure 1:**
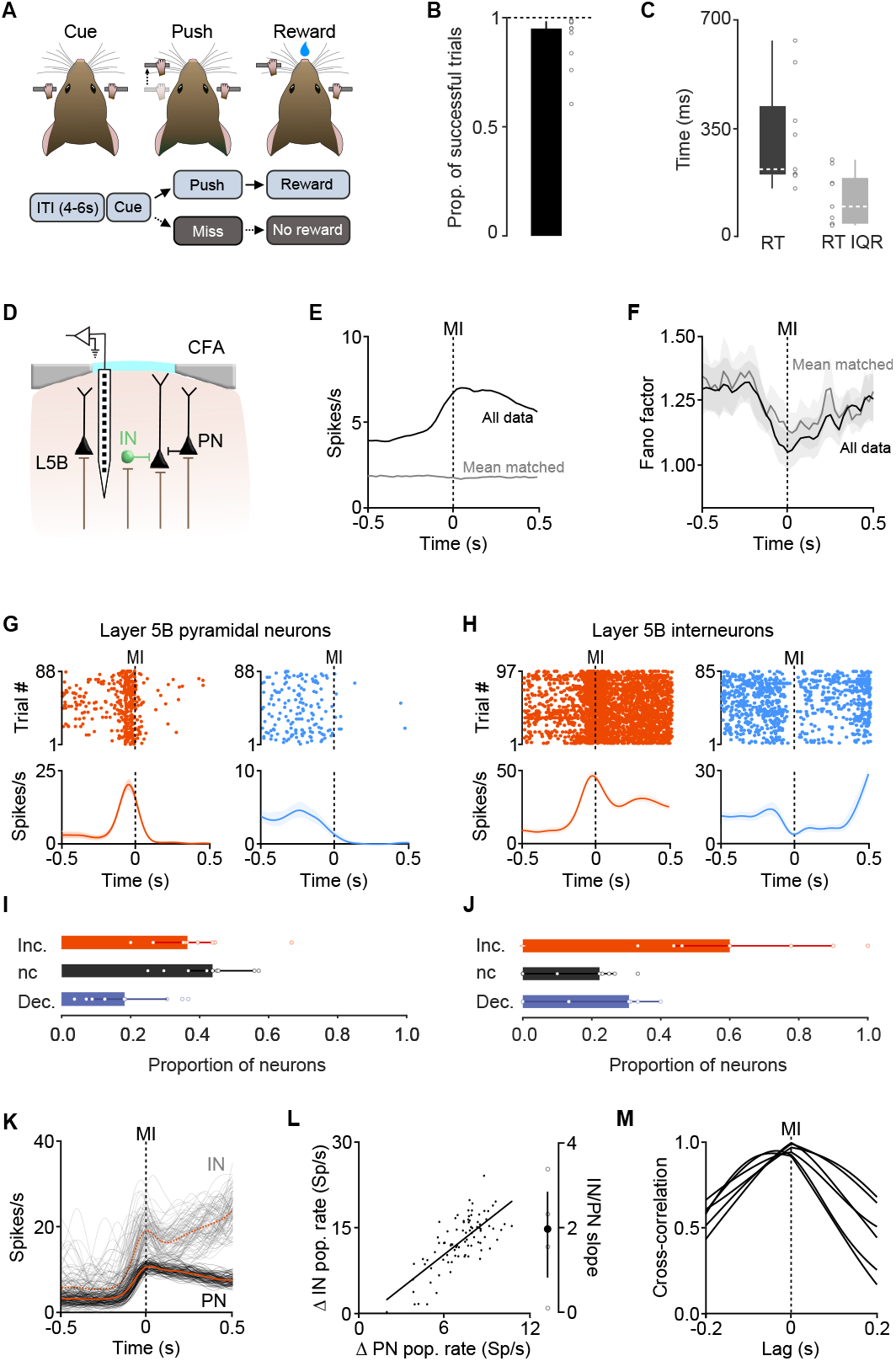
Co-activation of excitatory and inhibitory neurons align with neuronal variability quenching. (A) Top: cued forelimb push task. Bottom: behavioral task structure. ITI, inter-trial interval. (B) Task success as fraction of rewarded trials (Hit rate, N = 9 mice). Median +/-95% confidence intervals. Open circles, individual mice. (C) Median reaction times (RT, left) and variability of reaction times quantified as interquartile range (IQR, right). Dashed white lines, median. Open circles, individual mice. (D) High-density silicone probe recordings of putative layer 5B pyramidal neurons (PNs) and interneurons (INs) in the caudal forelimb area of motor cortex (CFA). (E-F) Change in population firing rate (E) or Fano factor (F) of layer 5B PN neurons aligned to movement initiation (MI) (n = 323 neurons, N = 9 mice). All data, black; mean matched data, gray. (G-H) Example response profiles of putative layer 5B pyramidal neurons (G) and interneurons (H) displaying increased (red) or decreased (blue) activity aligned to movement initiation (MI). Pyramidal neurons: increased activity, n = 117/323 neurons; decreased activity, n = 73/323 neurons. Interneurons: increased activity, n = 45/75 neurons; decreased activity, 15/75 neurons (N = 9 mice). Top, raster plots showing individual spike times across trials. Bottom, mean spike rate ± 95% confidence intervals. (I) Proportion of pyramidal neurons displaying increased (red), decreased (blue) or no change (nc) in activity during movement (N = 9 mice). Bars, median ± 95% confidence intervals; open circles, individual mice. (J) Proportion of interneurons displaying increased (red), decreased (blue) or no change (nc) in activity during movement (N = 9 mice). Bars, median ± 95% confidence intervals; open circles, individual mice. (K) Population firing rates for putative pyramidal neurons (black lines: single trials, solid red line: across-trial average) and interneurons (gray lines: single trials, dotted red line: across-trial average), aligned to movement initiation (MI) in an example session. (L) Left, population firing rate changes in putative pyramidal neurons and interneurons for an example session. Circles, individual trials; line, best fit to individual trial data. Right, median linear regression slope +/-95% confidence intervals across mice (N = 6 mice). (M) Cross-correlation of mean pyramidal neuron and interneuron population activity (N = 6 mice). Note, median delay = 0 ms.

Next, we compared the direction and timing of pyramidal neuron and interneuron activity changes (interneuron spike width <0.4 ms, n = 75 neurons from N = 9 mice) (Currie et al., 2022; Li et al., 2016) and found diverse response profiles with both neuron types showing increased and decreased activity, with a bias towards enhanced firing rates (Figures 1G-1J). Trial-by-trial population analysis revealed temporally correlated firing rate changes between pyramidal neurons and interneurons and co-varying magnitude (linear regression, p<0.05 in N = 5 / 6 mice, three mice excluded due to insufficient number of both neuron types, see Methods, median slope of firing rate change = 2.0 95% CI [0.8 2.9], median population lag = 0 ms, 95% CI [-23 0] (Figures 1K-1M and Figures S1A-S1C, N = 6 mice), suggestive of a common driving input.

The precise interneuron subtypes that contribute to quenching of neural variability remain unclear, but parvalbumin-(PV) and somatostatin-(SST) expressing populations have been suggested to play different roles in controlling network variability (Guo & Kumar, 2023; Haider et al., 2010; Jang et al., 2020; Onorato et al., 2025; Zhu et al., 2015). To investigate the activation of these interneuron subtypes, we performed cell-type-specific two-photon calcium imaging of layer 5 PV and SST interneurons during task engagement (Figure S1D). Consistent with our extracellular recordings, a substantial fraction of both interneuron subtypes were activated prior to movement initiation, concurrent with pyramidal neuron recruitment. However, their temporal dynamics diverged: PV interneuron activation was temporally aligned with that of pyramidal neurons, whereas SST interneurons exhibited delayed onsets. This temporal separation was evident both in the population-averaged activity (Figure S1E) and the distribution of single neuron onset times (Figures S1E-S1F), suggesting PV interneurons are the dominant source of local inhibition during movement initiation.

### Layer 5B pyramidal neurons receive coincident excitatory and inhibitory synaptic input during variability quenching

To investigate how the integration of excitatory and inhibitory synaptic inputs relates to variability quenching in single neurons, we performed whole-cell patch-clamp recordings in layer 5B pyramidal neurons (Figure 2A). These recordings enabled us to compare between single-cell membrane potential (*V_m_*) dynamics and the timing and magnitude of population-level variability quenching. We found that trial-by-trial *V_m_* variance mirrored changes in the population Fano factor with bidirectional changes in firing rate driven by either depolarizing (Δ*V_m_*, 3.3 mV 95% CI [2.1 8.4]) or hyperpolarizing (Δ*V_m_*, -2.0 mV 95% CI [-1.0 -4.0]) shifts in *V_m_* (Figure 2B-2D). Notably, the *V_m_* variance decreased uniformly across all neuronal response profiles, suggesting a common mechanism for variability quenching (Figure 2E). We next investigated the relationship between *V_m_* and variance during movement initiation and found that *V_m_* trajectories converged at a common membrane potential irrespective of the pre-movement baseline (Figure 2F and 2H). By comparing the trial-by-trial relationship between baseline *V_m_* and the movement-related shift in *V_m_* (Δ*V_m_*) we estimated the compound synaptic reversal potential (*E*_*rev*_) by extracting the voltage at which the predicted Δ*V_m_* equals zero from a linear fit of baseline *V_m_* and Δ*V_m_* (Figure 2G and 2I) (Sachidhanandam et al., 2013). Consistent with the direction of firing rate changes, *E*_*rev*_ was depolarizing in neurons that increased their firing rates (+3.3 mV 95% CI [2.4 8.0] relative to baseline *V_m_*) and hyperpolarizing in those that displayed decreased firing rates (-2.4 mV 95% CI [-0.9 -3.6] relative to baseline *V_m_*), with *E*_*rev*_ being close to the mean *V_m_* and hyperpolarized relative to spike threshold (*V*_*hresh*_) (Figure 2J and Figure S2A-S2C). These results suggest that individual motor cortex layer 5B pyramidal neurons receive a compound postsynaptic potential (PSP) during movement initiation that incorporates differing magnitudes of excitatory and inhibitory input (Sachidhanandam et al., 2013).

**Figure 2:**
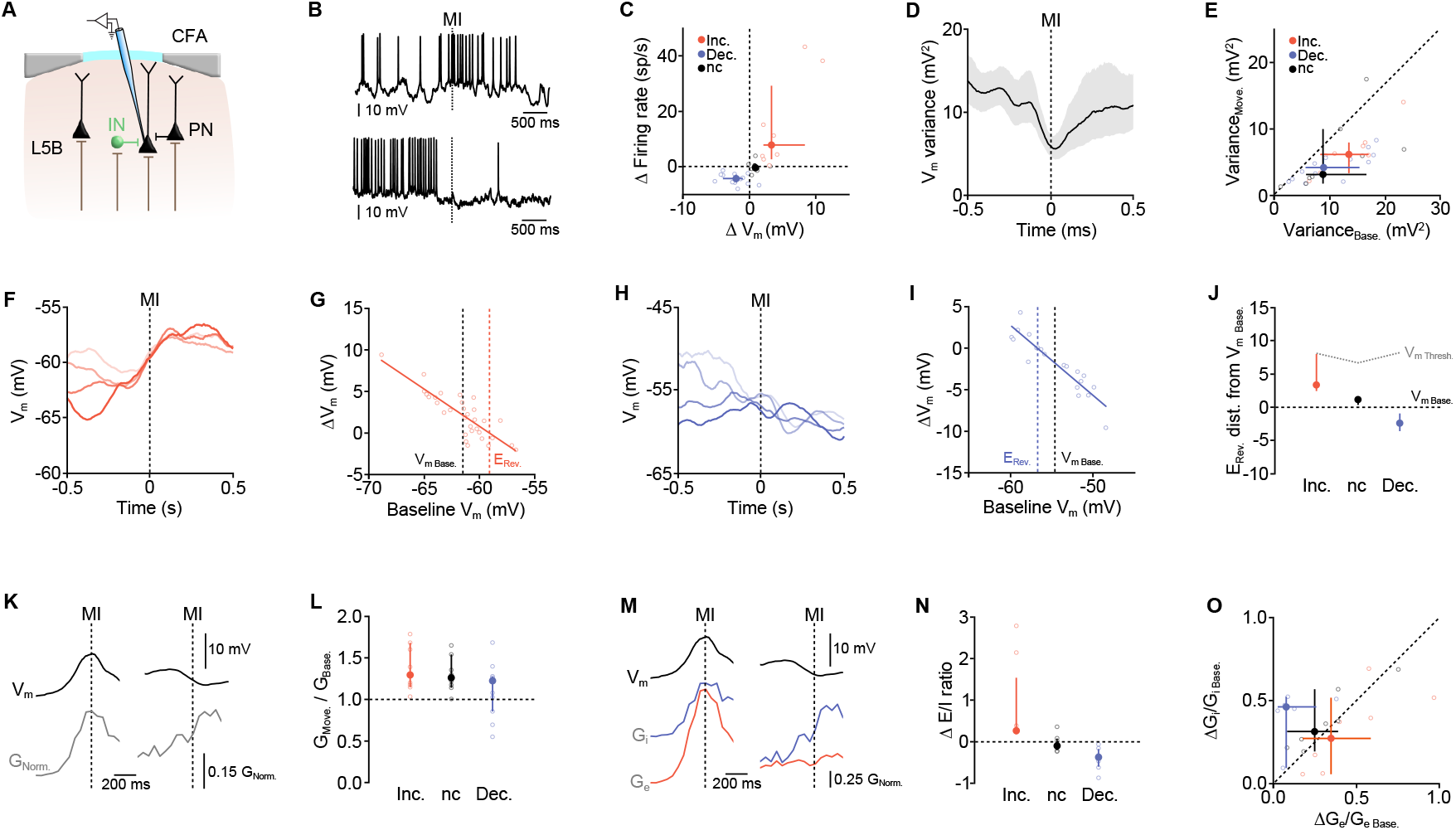
Coincident excitatory and inhibitory synaptic input drives neuronal variability quenching. (A) Whole-cell patch-clamp recordings from layer 5B pyramidal neurons (PNs) in the caudal forelimb region (CFA) of motor cortex. IN, interneuron; MTh, motor thalamus. (B) Membrane membrane potential recordings from layer 5B pyramidal neurons displaying increased (top) or decreased (bottom) firing rate during a lever push. MI, movement initiation. (C) Change in firing rate (spikes /s) as a function of subthreshold membrane potential in neurons with increasing (red), decreasing (blue) or no change (ns, black) in firing rate during movement initiation (n = 31 neurons, N = 31 mice). Filled circles, median ± 95% confidence interval (CI); open symbols, individual neurons. (D) Change in membrane potential (*V_m_*) variance of all pyramidal neurons (increased, decreased and no change) (n = 31 neurons, N = 31 mice). Black line, mean ± 95% CI; dashed line, movement initiation (MI). (E) Average *V_m_* variance during baseline (Base.) and movement (Move.) in layer 5B pyramidal neurons that display increased (red), decreased (blue) or no change in firing rate during movement initiation (n = 31 neurons, N = 31 mice). Filled circles, median ± 95% confidence interval (CI); open symbols, individual neurons, dashed line, unity. (F-G) *V_m_* trajectories from an individual pyramidal neuron displaying increased activity during movement initiation (MI). Traces are each averages of a quarter of trials sorted by baseline Base (F) and estimated synaptic reversal potential (*E*_*Rev*._, red dashed line) (G). *V*_*mBase*_ (black dashed line), mean baseline membrane potential. Thick line, best fit to individual trial data. (H-I) Base trajectories from an individual pyramidal neuron displaying decreased activity during movement initiation (MI) (H) and estimated synaptic reversal potential (*E*_*Rev*._, red dashed line) (I). *V*_*mBase*_ (black dashed line), mean baseline membrane potential. Thick line, best fit to individual trial data. (J) Synaptic reversal potential distance from baseline membrane potential ± 95% CI for pyramidal neurons that increase (red), decrease (blue) or show no change (nc, black) in firing rate during movement (n = 31 neurons, N = 31 mice). *V*_*mBase*_, normalised resting membrane potential; *V*_*mT hresh*._, spike threshold. (K) *V_m_* (top, black) and estimated compound conductance changes (bottom, gray) around movement initiation (MI). (L) Estimated peri-movement compound conductance (*G*_*Move*_) relative to baseline conductance (*G*_*Base*_)) in pyramidal neurons that increase (red), decrease (blue) or show no change (nc, black) in firing rate during movement (n = 27 neurons, N = 27 mice). Filled circles, median ± 95% confidence interval (CI); open symbols, individual neurons, dashed line, unity. (M) *V_m_* (top, black) and estimated excitatory (red) and inhibitory (blue) conductance changes (bottom) around movement initiation (MI). Note, 4 neurons were removed from the analysis due to negative conductance estimates. (N) Relative changes in the excitation-inhibition ratio (E/I ratio) for neurons that increase (red), decrease (blue) or show no change (nc, black) in firing rate during movement. Note, only neurons with a *G*_*Move*_/*G*_*Base*_ > 1 were included (n = 24/27 neurons, N = 24 mice). Filled circles, median ± 95% confidence interval (CI); open symbols, individual neurons, dashed line, unity. (O) Changes in estimated excitatory (*G*_*e*_) and inhibitory (*G*_*i*_) conductances relative to baseline conductance (*G*_*eBase*_ and *G*_*iBase*_) for neurons that increase (red), decrease (blue) or show no change (nc, black) in firing rate during movement. Note, only neurons with a *G*_*Move*_/*G*_*Base*_ > 1 were included (n = 24/27 neurons, N = 24 mice). Filled circles, median ± 95% confidence interval (CI); open symbols, individual neurons, dashed line, unity.

To estimate the underlying excitatory and inhibitory synaptic conductances, we exploited the fact that membrane capacitance and synaptic conductance kinetics are assumed to remain stable, such that conductances can be estimated from current-clamp data by analyzing the time constant of *V_m_* fluctuations (Berg & Ditlevsen, 2013). Applying this method, we found that most neurons displayed a net increase in conductance during movement initiation (n = 24 / 27 neurons from N = 27 mice; conductance ratio *G*_*Movement*_*/G*_*Baseline*_: increased 1.3 95% CI [1.1 1.7], decreased 1.2 95% CI [0.9 1.3], non-responsive 1.3 95% CI [1.1 1.5], Figures 2K-2L). While intrinsic voltage-dependent conductances could theoretically bias conductance estimates, a general bias is unlikely given the consistent conductance increase across neurons irrespective of response type (i.e. depolarizing / hyperpolarizing *V_m_* and increased / decreased spike rate) (see Figure 2C). We then decomposed the compound conductance into excitatory (*G*_*e*_) and inhibitory (*G*_*i*_) components by leveraging the distance of the *V_m_* from the theoretical excitatory (0 mV) and inhibitory (–75 mV) reversal potentials (Berg & Ditlevsen, 2013) (Figure 2M). Movement-related conductance changes and excitation/inhibition (E/I) ratios were consistent with the direction of *V_m_* and firing rate changes in individual neurons (Figure 2N). Increased firing rates were associated with coordinated increases in both *G*_*e*_ and *G*_*i*_ with a bias towards *G*_*e*_ (normalized Δ*G*_*e*_: 0.35 95% CI [0.18 0.59]; Δ*G*_*i*_: 0.27 95% CI [0.06 0.52], Figure 2O). In contrast, neurons with decreased firing rates displayed a preferential increase in *G*_*i*_ (normalized Δ*G*_*e*_: 0.08 95% CI [0.03 0.26]; normalized Δ*G*_*i*_: 0.46 95% CI [0.09 0.52], Figure 2O). As predicted, neurons that maintained baseline *V_m_* and firing rates during movement displayed balanced changes in *G*_*e*_ and *G*_*i*_ (normalized Δ*G*_*e*_: 0.25 95% CI [0.08 0.39]; normalized Δ*G*_*i*_ 0.31 [0.19 0.56], Figure 2O). Together, these results support a model whereby co-timed excitatory and inhibitory synaptic input and the resulting increase in compound conductance drive the membrane potential towards the reversal potential of that neuron, funneling the *V_m_* to an input-specific set-point (at *E*_*rev*_) to reduce both trial-by-trial membrane potential and firing rate variability.

### Compound synaptic input quenches membrane potential variability

To test this idea further, we developed a conductance-based point neuron model with excitatory (*G*_*e*_) and inhibitory (*G*_*i*_) synaptic conductances and baseline conductance (*G*_*Base*_) representing both intrinsic leak and background synaptic conductances (Figure 3A). Our in vivo recordings indicated only modest changes in mean *V_m_* at movement onset, with a depolarization of approximately 3 mV that could, in principle, arise from either relatively weak excitatory drive alone, or from stronger approximately balanced excitatory and inhibitory inputs, as implied by our synaptic reversal potential and conductance measurements. If both input structures can generate the same voltage change in individual neurons, what is the advantage of combined E/I inputs during movement initiation? We predicted that combined E/I input, scaled to produce the same net somatic depolarization as weak excitation alone, would suppress variability more effectively given that the higher total conductance would more efficiently drive the membrane potential toward its input-specific set-point (at *E*_*rev*_). We tested this prediction by modeling both scenarios and changing the initial membrane potential to mimic trial-by-trial pre-movement differences in the baseline E/I ratio. For simplicity and interpretability, we kept the background conductance constant across trials and omitted any fast in vivo - like fluctuations. We found that weak excitatory conductances generated a modest reduction in *V_m_* variance, while stronger combined E/I conductances generated significantly stronger variability quenching (Figures 3B-3C). Note that in both cases identical synaptic conductances were applied trial-by-trial, so the more effective quenching for the compound conductance is not due to less input variability. Consistent with the idea that increased synaptic conductance underlies variability quenching, our model also showed that strong excitatory drive alone can reduce trial-by-trial variability as effectively as a combined excitatory-inhibitory input, but at the cost of producing membrane depolarizations that fall outside the physiological range (i.e., generating single excitatory postsynaptic potentials of approximately +25 mV) (Figure S3). This suggests that combined excitatory-inhibitory input likely represent a canonical mechanism for minimizing *V_m_* and spike rate variability, while maintaining physiologically relevant voltage fluctuations. As the ratio of total synaptic conductance (combined *G*_*e*_ + *G*_*i*_ + *G*_*Base*_ = *G*_*Move*_) to baseline conductance (*G*_*Base*_) increased, membrane potential trajectories were driven towards the synaptic reversal potential more effectively, reducing movement-related trial-by-trial *V_m_* variability. This effect recapitulated estimated conductance changes and variability quenching observed in vivo (Figure 3D). We next investigated whether variability in the magnitude of inputs (*G*_*e*_ and *G*_*i*_) affected variability quenching and found that uncorrelated input variability, where trial-by-trial *G*_*e*_ and *G*_*i*_ magnitudes are variable and independent, increased *V_m_* variance, an effect that became more pronounced as conductance amplitude increased (Figure 3E). Conversely, correlated *G*_*e*_ and *G*_*i*_ input variability, where trial-by-trial *G*_*e*_ and *G*_*i*_ magnitudes are variable but co-modulated, reduced *V_m_* variance irrespective of the conductance magnitude (Figure 3F). These findings demonstrate that strong, co-varying excitatory and inhibitory synaptic input provides a biophysical mechanism to consistently drive individual neuron *V_m_* trajectories towards their combined synaptic reversal potential. This reduces trial-by-trial membrane potential fluctuations ensuring robust quenching of firing rate variability at the point of movement initiation.

**Figure 3:**
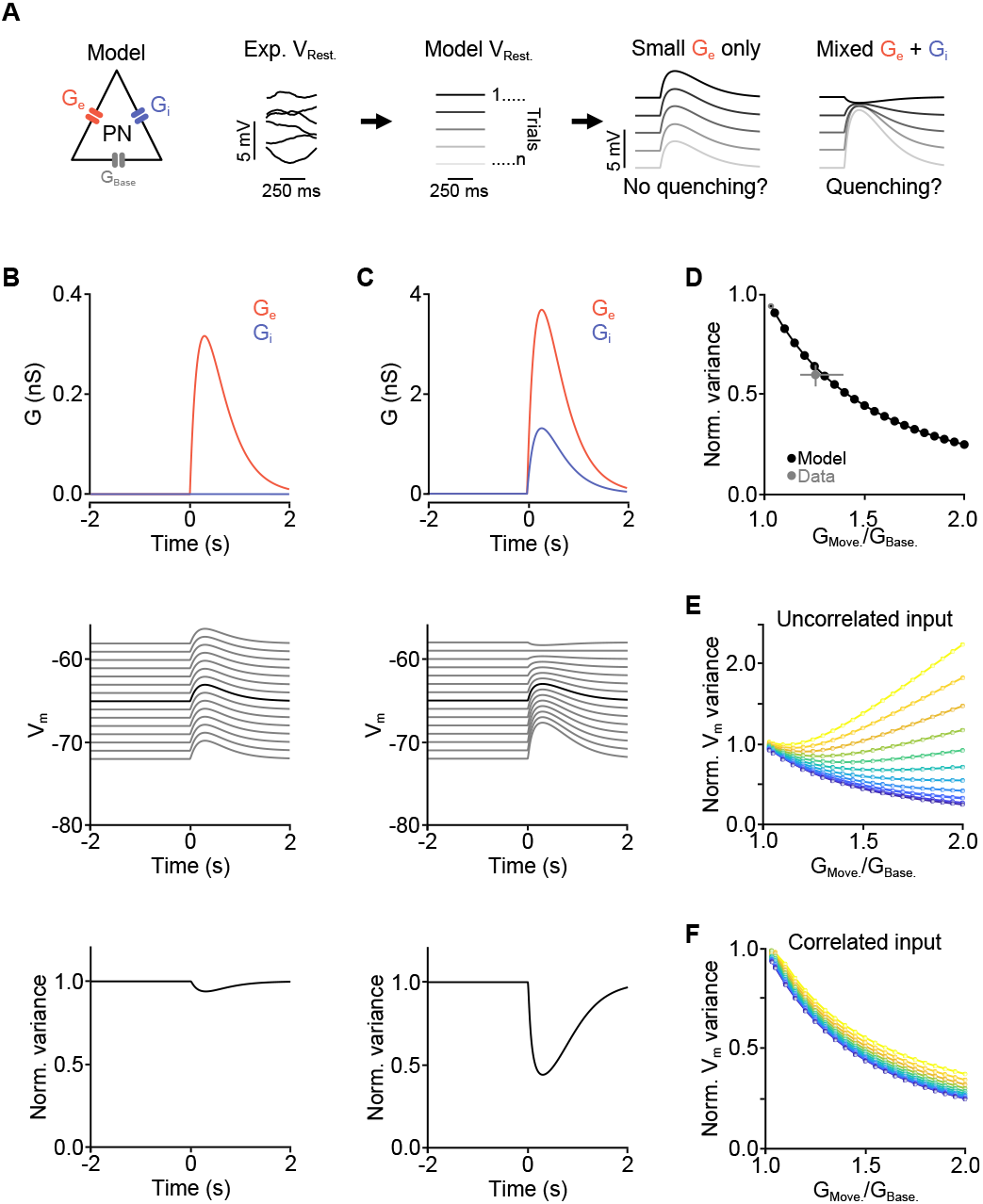
Large compound synaptic input drives *V_m_* variability quenching. (A) Conductance-based model including an excitatory (*G*_*e*_), inhibitory (*G*_*i*_) and baseline (*G*_*Base*_) conductance used to simulate effects of small excitatory or mixed excitatory and inhibitory inputs. Hypothesis to be tested - strong compound synaptic input drives quenching, whereas small inputs alone are insufficient to quench neuronal variability. Exp. *V*_*Rest*_., experimentally measured resting membrane potential; Model *V*_*Rest*_., simulated resting membrane potential to match in vivo recordings. (B) Simulated excitatory input alone (top) and resulting membrane potential changes (*V_m_*, middle) and normalized variance (bottom). (C) Simulated mixed excitatory and inhibitory input (top) and resulting membrane potential changes (*V_m_*, middle) and normalized variance (bottom). (D) Normalized variance as a function of the ratio of conductance during movement (*G*_*Move*_) and baseline (*G*_*Base*_). Black circles, model data; gray circle, median layer 5B pyramidal neuron data ± 95% confidence intervals (n = 24 neurons, N = 24 mice). (E-F) Effect of uncorrelated (E) versus correlated (F) input size of excitatory and inhibitory conductance on normalized variance. Input variability increases from 0 to +-100% for blue to yellow traces.

### Thalamic input quenches membrane potential variability during movement initiation

In rodents, thalamocortical input drives patterns of motor cortical activity necessary for movement (Dacre et al., 2021; Sauerbrei et al., 2020; Tanaka et al., 2018). To test whether feedforward input from the thalamus is necessary for neuronal variability quenching, we analyzed an existing dataset of intracellular recordings from layer 5B neurons in CFA during optogenetic stimulation of the ventral anterolateral (VAL) motor thalamus in the absence of an auditory cue (i.e. during the task inter-trial interval) (Figure 4A) (Dacre et al., 2021). Photostimulation of motor thalamus generated a robust increase in the compound synaptic conductance in most neurons and a reduction in *V_m_* variance comparable to that observed during cue-evoked movement initiation while also directly evoking lever pushes in the absence of a cue (photostimulation evoked movement in 71 ± 22% of trials, normalized post-photostimulation *V_m_* variance: 0.39 95% CI [0.25 0.50], normalized *V_m_* variance during cued movements 0.29 95% CI [0.21 0.52], Wilcoxon p = 0.56, N = 6) (Figures 4B-4E), indicating that thalamic input is sufficient to drive variability quenching. We then investigated whether motor thalamic input was necessary for variability quenching by comparing task related *V_m_* variability before and after thalamic infusion of the GABAA receptor agonist Muscimol (Figure 4F). Blocking thalamic input reduced task performance and *V_m_* variability quenching (pre Muscimol *V_m_* norm. variance 0.47, 95% CI [0.32 0.73], post Muscimol *V_m_* norm. variance 0.77, [0.66 0.83], N = 10, Wilcoxon p = 0.027; successful pushes pre Muscimol in 86 ± 18% of trials, successful pushes post Muscimol in 24±31% of trials) (Figure 4H) and abolished the task-related increase in synaptic conductance, indicating that thalamocortical input is a prerequisite for movement-related neuronal variability quenching (Figures 4G-4J).

**Figure 4:**
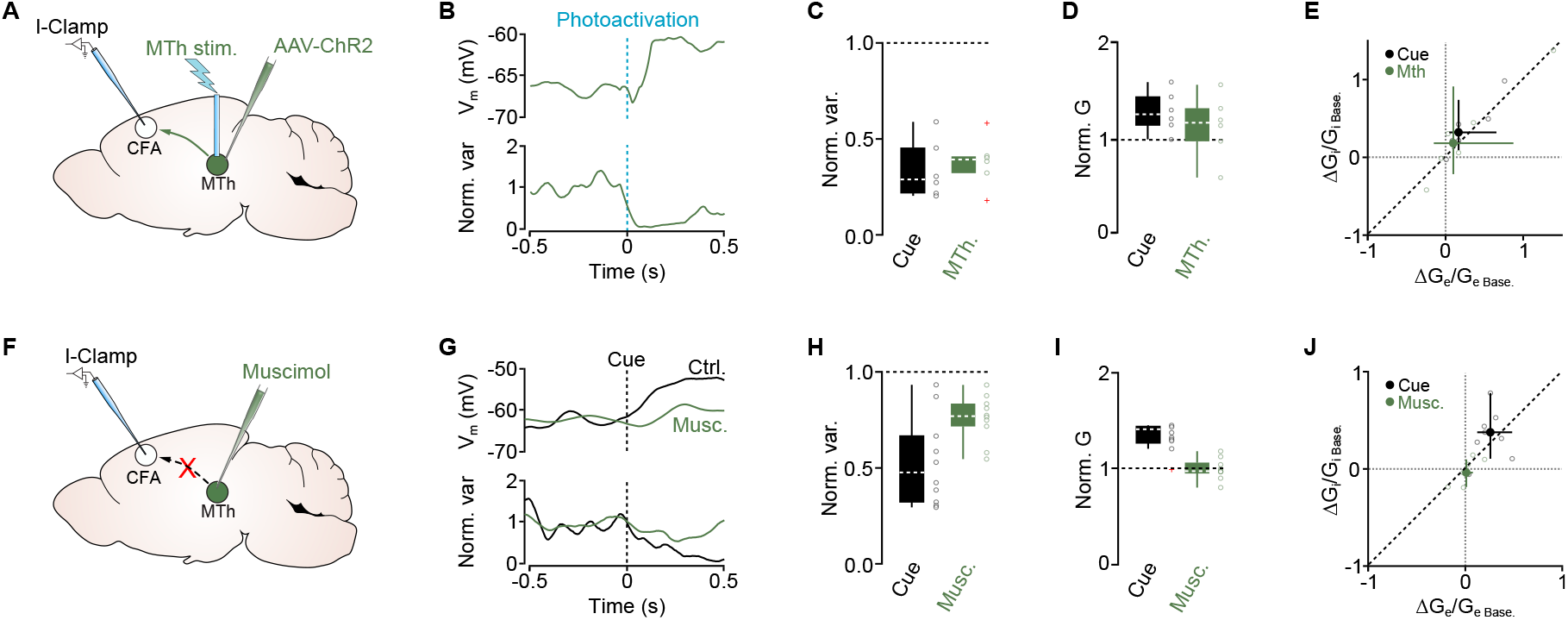
Thalamic input quenches *V_m_* variability during movement initiation. (A) Patch-clamp recordings in layer 5B CFA during photoactivation of motor thalamus (MTh). I-Clamp, current clamp. ChR2, Channelrhodopsin2. (B) Change in membrane potential (*V_m_*, top) and normalized variance (var. bottom) in a single pyramidal neuron during photoactivation of motor thalamus (MTh). Dashed line, photoactivation onset. (C-D) Box plots showing normalized variance (C) and conductance (D) after cue presentation (Cue) or photoactivation of motor thalamus (MTh) (n = 6 neurons, N = 6 mice,). Dashed white line, median; open circles, data from individual mice; red cross, outlier. (E) Normalized excitatory and inhibitory conductance (G) changes after cue presentation (Cue) or photoactivation of motor thalamus (MTh) (n = 6 neurons, N = 6 mice). Filled circles, median ± 95% confidence interval (CI); open circles, data from individual mice. (F) Patch-clamp recording in layer 5B CFA during Muscimol inactivation of motor thalamus (MTh). I-Clamp, current clamp. (G) Change in membrane potential (*V_m_*, top) and normalized variance (var. bottom) in a single pyramidal neuron during Muscimol inactivation of motor thalamus (MTh). (H-I) Box plots showing normalized *V_m_* variance (H) and conductance (I) after cue presentation (Cue) or Muscimol inactivation of motor thalamus (MTh) (n = 9 neurons, N = 9 mice). Dashed white line, median; open circles, data from individual mice; red cross, outlier. Note, 1 neuron was removed from the analysis due to negative conductance estimates. (J) Normalised excitatory and inhibitory conductance (G) changes after cue presentation (Cue) or Muscimol inactivation of motor thalamus (MTh) (n = 9 neurons, N = 9 mice). Filled circles, median ± 95% confidence interval (CI); open circles, data from individual mice.

### Thalamic input drives reliable trial-by-trial population dynamics in motor cortex

Our findings demonstrate that thalamic input alone can drive quenching of neuronal variability in single neurons, but how does this relate to changes in population dynamics and behavior? We leveraged the fact that optogenetic activation of motor thalamus directly, or via projections from the cerebellum, triggers lever pushes in the absence of a cue, but not in every trial (Figures 5A and 5B) (Dacre et al., 2021). This observation allowed us to compare population firing rate variability during cued or photostimulated push movements with trials where the same photostimulation failed to evoke a lever push (photostimulated miss trials). Across all conditions, the activity of pyramidal neuron and interneuron populations co-varied, indicating that thalamocortical input recruits excitatory and inhibitory subpopulations in a coordinated manner (Figures 5C-5E). The magnitude of population responses during photostimulated push trials was comparable to that of control (Figure 5F). However, in photostimulated miss trials, firing rate changes were significantly lower than those observed during push trials. Thus, thalamic stimulation can recapitulate cortical network dynamics associated with natural movement initiation, but the effect of photostimulation can differ from trial to trial.

**Figure 5:**
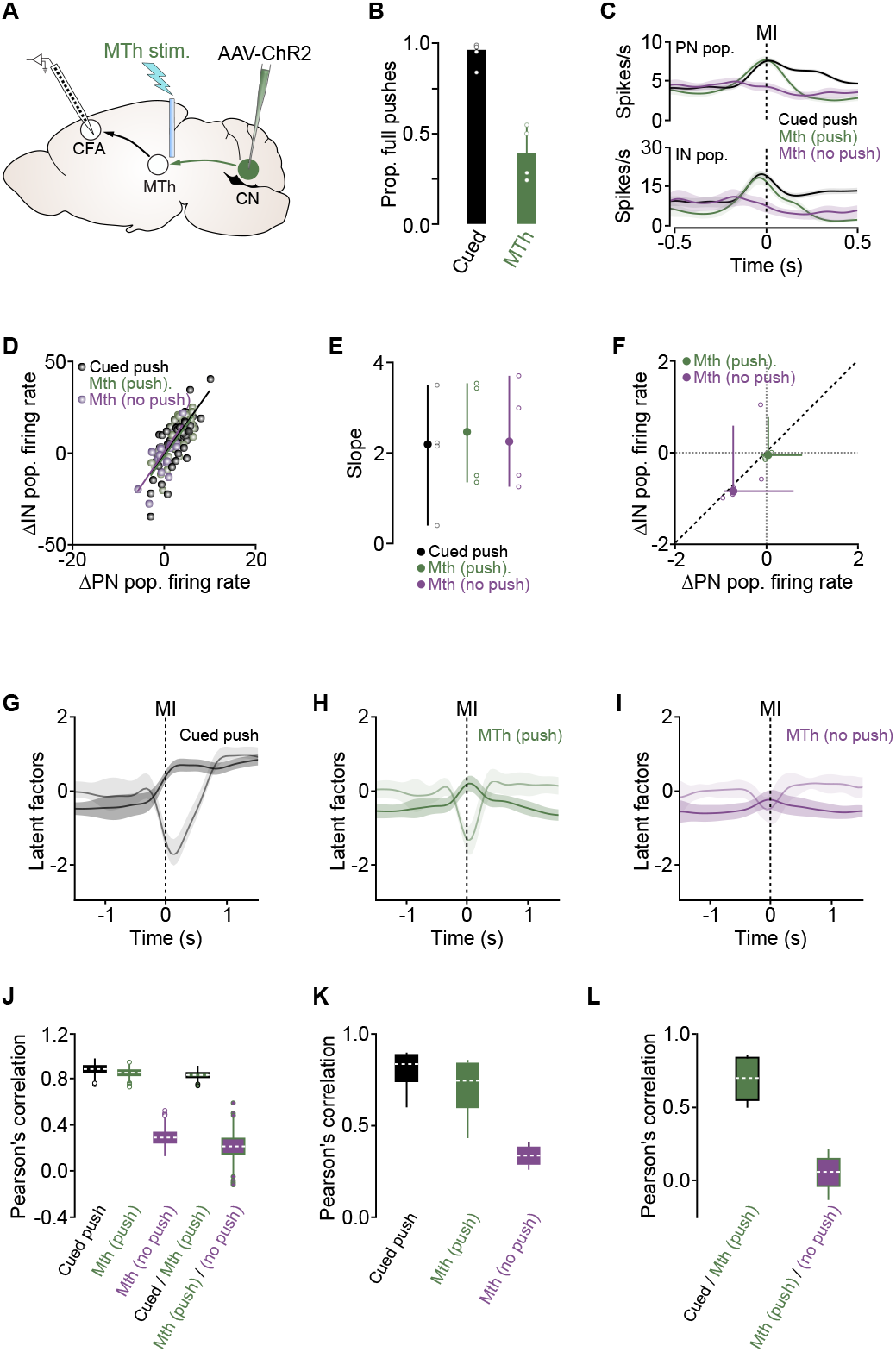
Thalamic input drives reliable trial-to-trial population dynamics during movement initiation. (A) High-density silicone probe recordings in CFA during optogenetic stimulation of motor thalamus. CFA, caudal forelimb area of motor cortex; MTh, motor thalamus; ChR2, channelrhodopsin2; CN, cerebellar nuclei. (B) Proportion of cued or photostimulation (MTh) trials with full lever pushes (N = 4 mice). Data, median +/-95%CI. Open circles, data from individual mice. (C) Change in firing rate of putative pyramidal neuron (PN pop., top) and interneuron population (IN pop., bottom) during cued trials (black), photostimulation trials with full lever pushes (MTh push, dark green) or photostimulation trials in the absence of lever pushes (MTh no push, light green). Data are from an example session within a single mouse. MI, movement initiation. (D) Population firing rate changes in putative pyramidal neurons and interneurons during cued trials (black), photostimulation trials with full lever pushes (MTh push, dark green) or photostimulation trials in the absence of lever pushes (MTh no push, light green). Data are from an example session within a single mouse. MI, movement initiation. Circles, data from individual trials; thick lines, best fit to individual trial data. (E) Average linear regression slope across mice for cued trials (black), photostimulation trials with full lever pushes (MTh push, dark green) or photostimulation trials in the absence of lever pushes (MTh no push, light green) (N = 4 mice). Open circles, data from individual mice. Closed circles, median +/-95%. (F) Changes in putative pyramidal neuron and interneuron population firing rates (MTh push & MTh no push) relative to population firing rate changes during cued trials (N = 4 mice). Open circles, data from individual mice. Closed circles, median +/-95%. (G-I) Gaussian Process Factor Analysis (GPFA) with a Gaussian emission model was used to obtain low-dimensional representations of population activity. Population activity projected onto the first two dimensions (latent factor 1, dark color; latent factor 2, light color) during cued trials (black), photostimulation trials with full lever pushes (MTh push, dark green) or photostimulation trials in the absence of lever pushes (MTh no push, light green). MI, movement initiation. (J) Bootstrapped trial-wise Pearson correlation between latent factor trajectories of the example mouse in (G)-(I) comparing trials within the same trial type (cued push, MTh push, MTh no push) and across trial types (cued/MTh push, Mth push/ Mth no push). Dashed white lines, median. (K) Within-trial-type correlation (bootstrapped) of latent factor trajectories across N = 4 mice. Dashed white lines, median. (L) Across-trial-type correlation (bootstrapped) of latent factor trajectories across N = 4 mice. Dashed white lines, median.

To assess the relationship between the variability in motor cortical population dynamics and behavior more directly, we projected the population activity onto a low-dimensional latent space to compare the consistency of trial-by-trial trajectories across different trial types (Figure 5G-5I). Both cued and photostimulated push trials exhibited high across trial correlations in their latent factors within and across conditions, indicative of low variability and similar population dynamics during movement initiation. Differences in cued trial latent factor dynamics post movement (i.e. >200ms after movement initiation) likely reflect reward anticipation and consumption, which were unique to this trial type. In contrast, photostimulated miss trials displayed significantly lower across trial correlations and weaker across condition similarity (Figures 5J-5L, Cued push vs photostimulation miss p = 0.003, photostimulation push vs photostimulation miss, p = 0.03), indicating highly variable trial-by-trial population responses. This suggests that long-range feedforward thalamic input drives robust variability quenching and reliable, structured trial-by-trial population dynamics during movement initiation.

## Discussion

By recording sub- and supra-threshold activity in primary motor cortex during cued forelimb lever movements, we discovered that trial-by-trial firing rate variability is quenched at movement initiation via a synaptic funneling mechanism, driven by thalamocortical input. Co-timed excitation and inhibition produce a profound increase in total synaptic conductance, driving layer 5B pyramidal neurons towards their compound synaptic reversal potential that stabilizes neuronal output by reducing *V_m_* and firing rate variability.

Although the increased synaptic conductance during movement initiation could originate from diverse sources (Goz & Hooks, 2023; Hooks et al., 2013; Kristl et al., 2025; Luo et al., 2019; Okoro et al., 2022; Zingg et al., 2014), we identified motor thalamus as a critical driver. Thalamic input was both necessary and sufficient for variability quenching, a conclusion supported by gain- and loss-of-function manipulations that validate recent theoretical predictions regarding the role of external input in driving variability quenching (Goris et al., 2024). While we previously established that coordinated input from the ventral anterolateral motor thalamus gates cortical output for movement initiation (Dacre et al., 2021), our current findings reveal the biophysical substrate through which this feedforward circuit ensures robust and reproducible quenching at movement initiation. Although thalamus directly targets layer 5B pyramidal neurons, local recurrent networks will also contribute (Wu et al., 2025). For example, PV interneurons, readily activated by single synaptic inputs (Jouhanneau et al., 2018), can be recruited via feedforward and recurrent pathways. Contrary to models of output gating via disinhibition, we found that PV-mediated inhibition increased in parallel with excitation, helping to define a neuron-specific synaptic reversal potential (Estebanez et al., 2017; Isomura et al., 2009; Kaufman et al., 2013). Critically, variability quenching was observed across neurons regardless of whether net firing rates increased or decreased, implying an input-driven balanced E/I regime that stabilizes motor output across trials (Guo & Kumar, 2023; Hennequin et al., 2018; Rostami et al., 2024; van Vreeswijk & Sompolinsky, 1996). In theory, unopposed feedforward excitation would also quench variability via increased conductance, but at the cost of runaway activity that would eventually saturate the firing rate dynamic range (Chance et al., 2002). Instead, the motor cortex orchestrates correlated and balanced shifts in excitation and inhibition in the network. This configuration produces the conductance increase necessary for quenching while permitting modest scaling of output firing rates, thereby ensuring both stable and reproducible network dynamics during motor output.

To what extent does the observed single neuron mechanism scale to population-level responses? We show that thalamically-driven, conductance-based quenching at the single-neuron level is both necessary and sufficient for generating reliable population dynamics. Our population analyses revealed that successful movement initiation, whether cued or optogenetically triggered, is characterized by low trial-by-trial variability and highly stereotyped latent dynamics within the motor cortical population. In contrast, failure to stimulate movement (photostimulated miss trials) was associated with a lack of quenching and unreliable population activity. The reason for this difference remains unclear but could reflect differences in thalamic and cortical network state caused by dynamic switching of attention, arousal, or behavioral states that gate output-potent cortical configurations (Dacre et al., 2021; Halassa et al., 2014; Kaufman et al., 2014; Poulet et al., 2012; Poulet & Petersen, 2008; Schiemann et al., 2015). The level of synchrony in cortical circuits, determined by the modulation of brain state, may be a determinant factor permitting the coordinated recruitment of excitation and inhibition upon thalamic input in motor cortex (McCormick et al., 2020; Pattadkal et al., 2025; Tan et al., 2014). In this regard, thalamus would provide the essential trigger, but the resultant quenching may also reflect a positive feedback loop where stabilized single-neuron responses facilitate the emergence of stable attractor states at the population level (Hennequin et al., 2018).

We suggest that this simple, conductance based biophysical mechanism for variability quenching may generalize across sensorimotor systems. Variability quenching is a widespread cortical phenomenon, prominent during sensory processing and task engagement. In mouse sensory areas, including somatosensory (Sachidhanandam et al., 2013) auditory (Kato et al., 2017), and visual (Adesnik, 2017) cortices, stimulus onset evokes a substantial increase in synaptic conductance. Similarly, in mouse premotor cortex, a task-related reduction in subthreshold variance is linked to faster membrane potential fluctuations, a hallmark of elevated conductance (Inagaki et al., 2019). These collective observations suggest that coordinated excitatory and inhibitory drive, resulting in high synaptic conductance, could be a conserved mechanisms for stabilizing cortical activity. However, this dominant conductance-based mechanism likely operates in parallel with other ways to reduce firing rate variability (Pattadkal et al., 2025). For example, hyperpolarized membrane potential responses can result from a net reduction in synaptic conductance, driven by interneuron-mediated network suppression (Aponte et al., 2021; Kato et al., 2017). In our study, a small subset of neurons (3 of 27) exhibited this pattern of activity (reduced conductance alongside decreased variance) indicative of parallel cooperative mechanisms for variability quenching within the same cortical area.

While our study identifies a simple feedforward mechanism for variability quenching, several fundamental questions remain. How do different behavioral states such as arousal, attention or motivation enable thalamic input to generate cortical activity patterns permissive for movement initiation (Dacre et al., 2021; Dura-Bernal et al., 2023; Inagaki et al., 2022; Kaufman et al., 2014; McCormick et al., 2020)? Does this conductance-based quenching mechanism in fact generalize to other cortical areas and behaviors or are there area-specific biophysical differences related to the underlying computations being performed? To what extent does pathological disruption of this synaptic funneling mechanism, through altered thalamocortical connectivity or E-I balance, contribute to atypical neural dynamics observed in movement disorders? Addressing these questions will bridge cellular mechanisms to systems-level computations and reveal how the brain dynamically tunes neural variability to balance stability and reproducibility with flexibility during behavior.

## Acknowledgements

We would like to thank Janelle Pakan for technical assistance with 2-photon imaging and discussions, Ayisha Mahmood, Thomas Clarke and Marie Zechner for assistance in training mice and Jessica Passlack, Constantinos Eleftheriou and Michelle Sanchez Rivera for comments on the manuscript. The research was supported by grants from the Biotechnology and Biological Sciences Research Council (BB/Y004639/1. to I.D.., BB/X01861X/1 to MH), a Wellcome Senior Research Fellowship and Wellcome Discovery Award (110131/Z/15/Z & 227475/Z/23/Z to I.D.), the German Research Foundation (DFG, AM 443/1-1 and AM 443/3-1 (539485808) to J.J.A), the Simons Initiative for the Developing Brain (to N.L.R. & I.D.), the Wellcome Trust and the Royal Society (Sir Henry Dale fellowship to N.L.R.), the European Research Council under the European Union’s Horizon 2020 research and innovation program (grant agreement 866386 to N.L.R.), the European Molecular Biology Organisation (YIP award to N.L.R.)

## Methods

### Mice and husbandry

All experiments and procedures were approved by the University of Edinburgh local ethical review committee and performed under license from the UK Home Office in accordance with the Animal (Scientific Procedures) Act 1986. Male adult C57BL/6J wild-type (RRID: IMSR_JAX:000664), PVCre Pvalb<tm1(cre)Arbr>) (RRID:IMSR_JAX:017320) or SST-Cre (Sst<tm2.1(cre)Zjh>) RRID:IMSR_JAX:013044 (8-14 weeks old, 20-35 g, one to six animals per cage) were maintained on a reversed 12:12 hour light:dark cycle (lights off at 7:00 am) and provided ad libitum access to food and water except during behavioral training and experimentation (see below).

### General surgery

Mice were induced under 4% and maintained under ∼ 1.5% isoflurane anesthesia, with all animals receiving a fluid replacement therapy (0.5 ml sterile Ringer’s solution; to maintain fluid balance) and buprenorphine (0.5 mg/kg; for pain relief) or buprenorphine (0.5 mg/kg) and carprofen (0.5 mg/kg) at the beginning of each surgery. A lightweight headplate (∼ 0.75 g) was implanted on the skull using cyanoacrylate super glue and dental cement (Lang Dental, USA). Craniotomies were performed in a stereotactic frame (Kopf, USA) using a hand-held dentist drill with 0.5 mm burr (craniotomy diameter: whole-cell patch-clamp recording or viral injection ∼ 300 -1000 *µm*, cranial window surgery 2-4 mm. Viral vectors were delivered via pulled glass pipettes (5 *µl*, Drummond) using an automated injection system (Model Picospritzer iii, Intracell). After surgery, mice were left for 24-72 hours to recover. For postoperative care either buprenorphine (0.5 mg/kg) was administered in the form of an edible jelly cube ∼24 h after recovery from surgery or carprofen was injected for 72 h post-surgery every 24 h. At the end of each experiment, mice were anesthetized with euthatal (0.10–0.15 ml) and transcardially perfused with 30 ml of ice-cold 0.1 M phosphate-buffered saline (PBS) followed by 30 ml of 4% paraformaldehyde (PFA) in 0.1 M PBS solution. Brains were post-fixed in PFA overnight at 4°C then transferred to 10% sucrose solution for longer-term storage.

### Behavioral training

Mice were handled extensively before being head restrained and habituated to the behavioral setup. Mice were placed on a water control paradigm (1 ml/day) and weighed daily to ensure body weight stayed above 85% of baseline weight. Mice were trained for 30-minutes per day and had to wait during a random inter-trial-interval (ITI) of 4-6 s, before pushing the lever 4 mm forward in response to a 6 KHz auditory go cue to receive a ∼5 *µl* water reward. The duration of the auditory cue and the response window was shortened in progressive training stages: stage 1 = 10 s, stage 2 = 5 s, stage 3 = 2 s, with mice progressing to the next stage of training after achieving > 70 rewards in a single session or > 50 rewards in two consecutive sessions. Premature lever movements during the ITI resulted in a lever reset and start of a new ITI. After each session, mice were removed from the head restraint and given the remainder of their daily water allowance before being returned to their home cage.

### Extracellular recording

Single unit recordings during lever push trials were reanalyzed from previously published datasets (Currie et al., 2022; Dacre et al., 2021). Briefly, extracellular unit recordings from CFA were acquired using acutely implanted silicone probes (Neuropixels Phase 3B probes, IMEC). Data were acquired from the 384 channels closest to the probe tip using SpikeGLX software (30 KHz, 500 amplifier gain and high-pass filtered with a cutoff frequency of 300 Hz). Probe location was confirmed via DiI (Thermofisher) reconstruction of the recording probe tract, and units from 500-1200 *µm* below the pial surface (i.e. putative layer 5) were included in the analyses.

### Whole-cell patch-clamp recordings

Whole-cell patch-clamp recordings targeted to layer 5B, 550–1000 *µm* from the pial surface, were obtained from awake, head-restrained mice after performing a craniotomy and durotomy centered above CFA. Signals were acquired at 20 kHz using a Multiclamp 700B amplifier (Molecular Devices) and low pass filtered at 10 kHz using PClamp 10 software in conjunction with a DigiData 1440 DAC interface (Molecular Devices). No holding current was injected during recordings and the membrane potential was not corrected for liquid junction potential. Patch pipettes (5.5-7.5 *M* Ω) were filled with internal solution (285-295 mOsm) containing: 135 K-gluconate, 7 KCl, 10 HEPES, 10 sodium phosphocreatine, 2 MgATP, 2 Na_2_ATP and 0.5 Na_2_GTP (pH 7.2, 285-295mOsm), and 1-2mg ml^*−*1^ biocytin was added immediately before recording. External solution contained: 150 mM NaCl, 2.5 mM KCl, 10 mM HEPES, 1 mM CaCl_2_, and 1 mM MgCl_2_ (adjusted to pH 7.3 with NaOH). 14 of the 31 neurons reported in Figure 1 and all 16 neurons reported in Figure 3 were re-analyzed from Dacre et al., 2021.

### 2-photon imaging

To perform population calcium imaging in layer 5B, we expressed GCaMP6s in pyramidal neurons and PV and SST interneurons. For interneuron specific imaging, we injected 200 nl of the adeno-associated virus (AAV) AAV1.Syn.Flex.GCaMP6s.WPRE.SV40 (Chen et al., 2013) or AAV1.CAG.Flex.Ruby2-GSG-P2A-GCaMP6s.WPRE-Pa (Rose et al., 2016) into CFA to label SST and PV interneurons in Cre-driver transgenic mice. For pyramidal neuron imaging, we injected AAV1.CaMKII0.4.Cre.SV40 (University of Pennsylvania Vector Core, PA) together with AAV1.Syn.Flex.GCaMP6s.WPRE.SV40 in C57Bl/6 wild type mice. For all injections we used a 45 degree angle to target layer 5B of the CFA of motor cortex (AP: 0.6, ML: 1.6, DV: 0.6 mm) using a pulled glass pipette (5 ml, Drummond Scientific; 20–40 nl/min) and automated injection system (Model Picospritzer iii, Parker), before sealing the craniotomy with silicone (Body Double; Smooth-On,PA, USA) and implanting a lightweight headplate. AAV1.Syn.Flex.GCaMP6s.WPRE.SV40 was a gift from Douglas Kim & GENIE Project (Addgene viral prep 100845-AAV1; http://n2t.net/addgene:100845; RRID:Addgene_100845).

pAAV-CAG-Flex-mRuby2-GSG-P2A-GCaMP6s-WPRE-pA was a gift from Tobias Bonhoeffer & Mark Huebener & Tobias Rose (Addgene viral prep 68717-AAV1; http://n2t.net/addgene:68717 ; RRID:Addgene_68717). For imaging, a cranial window (glass coverslip 0; Menzel-Gläser, Germany), was implanted above the virus injection site and secured with cyanoacrylate glue. 2-photon calcium imaging was performed using a custom-built resonant scanning 2-photon microscope (320 x 320 *µm* FOV; 600 x 600 pixels) with a Ti:Sapphire pulsed laser (Chameleon Vision-S, Coherent, CA, USA; < 75 fs pulse width, 80 MHz repetition rate) tuned to 920 nm wavelength. Images were acquired at 40 Hz with a 40x objective lens (0.8 NA; Nikon) and LabVIEW-based software (LotoScan). To facilitate reliable imaging of layer 5 neurons at depths > 500 *µm*, imaging was performed from 24 hrs post-surgery. The combination of low pixel dwell time and blanking of FOV edges, where the dwell time is higher, and the addition of room temperature artificial cerebrospinal fluid on the surface of the skull reduced the risk of thermal effects (as discussed in Currie et al., 2022; Podgorski & Ranganathan, 2016).

### Motor thalamic activation

For optogenetic activation of motor thalamus as described in Dacre et al., 2021, 250 nl of AAV1-CAG-ChR2-Venus (2.3×10^12^ GC/ml, Addgene 20071) (Petreanu et al., 2009) was injected into contralateral motor thalamus (AP: 1.1, ML: 1.0, DV: 3.4 mm) and stimulated via an optic fiber (200 *µm* diameter, 0.39 NA; Thorlabs, sealed with RelyX Unicem2 Automix cement, 3M)) implanted into contralateral thalamus (AP: 1.1, ML: 1.0, DV: 3.2 mm). pACAGW-ChR2-Venus-AAV was a gift from Karel Svoboda (Addgene viral prep 20071-AAV1); http://n2t.net/addgene:20071 ; RRID:Addgene_20071). Light with 473nm wavelength was delivered in pulse trains (5-8 mW, 16.6 Hz pulse frequency, 33.3% duty cycle) using a solid-state laser (DPSS, Civillaser, China) and shutter (LS3S2T1, Uniblitz) controlled by an Arduino control system coupled to the implanted optic fiber by an optic patch cable (Thorlabs, FT200UMT). For activation of motor thalamus via dentate/interpositus nucleus axon terminals, AAV1-CAG-ChR2-Venus (2.3×10^12^ GC/ml, Addgene 20071) was injected into ipsilateral dentate (AP: 6.0, ML: 2.25, DV: 2.6& 2.2 mm) and interpositus (AP: -6.0, ML: -1.75, DV: -2.4 mm) cerebellar nuclei, with 75 nl injected at each depth within each nucleus. The placement of the optic fibers and the stimulus protocols were identical to those described for direct motor thalamic stimulation. Before optogenetic stimulation experiments were performed, mice were trained and habituated to 473 nm masking light and shutter activation. During habituation and experimental sessions, mice were exposed to 3 different trial types: (1) cue and shutter (2) laser and shutter; and (3) shutter only. Trials were presented in the sequence: 1, 1, 3, 1, 1, 2, 1, 1, 3…repeating for 30 minutes. For histological confirmation of the injection site and optic fiber placement, mice were transcardially perfused, decapitated and the whole head (including headplate and optic fiber) was post-fixed in 4% PFA for 2 days to improve preservation of the optic fiber tract. Coronal sections (60 *µm*) of CFA and MThDN/IPN were cut with a vibratome, mounted with Vectashield, and imaged using a slidescanner (Axioscan, Zeiss). The center of the optic fiber (COF) was defined as the most ventral extent of the optic fiber tract across all slices from each brain as measured from the pial surface. Where tracts of equal depth were present, the coronal section containing the largest diameter tract tip was identified as the COF. The expression of ChR2-Venus in MThDN/IPN was coarsely defined by first referencing three coronal slices (120 *µm* spacing) centered on the COF to the Paxinos and Franklin Mouse Brain Atlas before manually evaluating the proportion of each of the principle motor thalamic nuclei (AM, anteromedial; VL, ventrolateral; VPM, ventral posteromedial nucleus; VPL, ventral posterolateral; VM, ventromedial) containing fluorescence, and categorizing three levels based on expression covering 0%–5%, 5%–50% and 50%–100% of each nucleus. Data were not included from mice in which the COF was misaligned to virus expression.

### In vivo pharmacology

As described in (Dacre et al., 2021), for thalamic inactivation during patch-clamp recordings, a second craniotomy was performed above motor thalamus under general anesthesia (AP: 1.1, ML: 1.0). After 5/10 minutes of baseline task execution, the lever was locked and a small volume of the GABAA receptor agonist Muscimol (dissolved in external solution containing 150 mM NaCl, 2.5 mM KCl, 10 mM HEPES, 1.5 mM CaCl_2_ and 1 mM MgCl_2_) or external solution was injected with a pipette into motor thalamus (200 nl of 1 mM Muscimol, AP: 1.1, ML: 1.0, DV: 3.4 mm) at a rate of 5-20 nl/s. To confirm the anatomical location of drug injection, 1% w/v of retrobeads (Lumaflor Inc.) was included in the Muscimol/saline solution and the locations were confirmed post hoc. Mice were randomly assigned to drug or control groups, and experimenters were blinded. After each experiment, mice were transcardially perfused and data from animals in which retrobeads were found within motor thalamus were analyzed.

## Analysis

### Extracellular data analysis

Spike sorting was performed using Kilosort3 to automatically cluster units from raw data (Pachitariu et al., 2016). The resulting spike clusters were manually curated using Phy (https://github.com/cortex-lab/phy), and any unit with sufficient refractory period violations, inconsistent waveform amplitude across the duration of the recording, or clipped amplitude distribution was excluded from analyses. In addition, we performed a variance ratio test for random walk on the spike count per trial to exclude neurons that showed substantial firing rate drift within a session, presumed to be due to subtle electrode movement. Neurons were defined as putative fast-spiking (FS) interneurons and putative regular spiking (RS) pyramidal cells according to their spike width, (spike width, FS < 0.4 ms; RS >= 0.4 ms). To restrict analysis to putative layer 5 neurons, we included single units between 500-1200 *µm* from pial surface. To detect changes in activity, firing rates were calculated by convolving spike times with a Gaussian kernel of 50 ms and neurons were classified as increasing or decreasing their activity based on their net changes between a window -500 - -400 ms prior to movement and a peri-movement period of -50 to +50 ms with a Wilcoxon signed rank test. We chose these time windows to be consistent with the analysis of changes in firing rate variability in movement aligned data (fano factor, see next paragraph.) Population Fano Factors were calculated using analysis code from (Churchland et al., 2010) (available from https://churchland.zuckermaninstitute.columbia.edu/content/code). We calculated the population Fano factor in a moving window of 100 ms with 25 ms time steps.

For comparison of baseline and peri-movement values we used the window at t = -500 ms relative to movement as baseline and t = 0 as the peri-movement window. The mean matched Fano factor was computed as in (Churchland et al., 2010). This procedure ensures that there are no systematic differences in firing rates between time bins which could bias the result given the variance depends on the mean firing rate. Briefly, mean matching is achieved by computing the distribu¬tion of mean counts across neurons for each time bin and calculating the greatest common distribution present at all times. For each time bin, the distribution of mean rates is then matched to this common distribution by randomly discarding data points until the distributions match before computing the Fano factor on the matched distribution. The mean matched Fano factor is then the mean of 50 repetitions of this procedure. The delay between excitatory and inhibitory subpopulations was calculated from the lag of the peak cross-correlation between the grand averages of the excitatory and inhibitory populations (signal correlation). For this analysis only sessions with at least 5 excitatory and 5 inhibitory neurons were used (N = 6/9 mice). To analyze trial-by-trial changes in excitatory and inhibitory populations we calculated the averages of the excitatory and inhibitory populations in each trial, subtracted the firing rate during the baseline window from the firing rate in the peri-movement window for these population averages in each trial and fit a linear regression model to the data across all trials (firing rate changes in inhibitory population vs firing rate changes in excitatory population). Gaussian Process Factor Analysis (GPFA) with a Gaussian emission model was used to obtain low-dimensional representations of the extracellularly recorded population activity (Yu et al., 2009). Combining Gaussian processes with factor analysis, GPFA uses moment representations to describe the evolution of latent factors driving observed neural dynamics (Hurwitz et al., 2021). Spike counts were binned at 50 ms resolution in a -2s to 2s window around movement onset. For each animal, a single model was fit to all combined trials in the cued condition and during motor thalamic activation. To align trials with motor thalamic activation without movement, we used the average reaction time from stimulation trials with movement. GPFA models were fitted using the implementation in the Elephant framework (Yegenoglu et al., 2018). We tested models with between two and 9 factors and observed a stabilization of the test likelihood with 8 factors. To aid visualization, the results presented in Figure 5 were obtained using the two factor models, and the trends were the same for the larger models.

### Whole-cell data analysis

All whole-cell recordings were analyzed using custom-written scripts in MATLAB. Individual action potentials (APs) were initially detected with a wavelet-based algorithm (Nenadic & Burdick, 2005) and false positives were removed with the help of semi-automated k-means clustering under visual inspection. Average AP firing frequencies were calculated by convolving spike times with a 50 ms Gaussian kernel. Significant modulation was tested comparing the firing rate during a baseline window at time = -400- -300 ms prior to movement with the firing rate in a peri-movement window of -50 to +50 ms relative to movement initiation with a Wilcoxon signed rank test and response types of individual neurons (increasing or decreasing firing rate at movement initiation) were based on their average firing rate changes between these time windows. For subthreshold trajectory analysis, APs were removed and the membrane potential filtered with a Savitzky Golay filter (order = 1, framelength = 2001). Slow baseline drifts during the recordings were removed by first calculating a median value from the *V_m_* distribution of the first 60s after equilibration and then subtracting the quantified drift (i.e. deviation from initial median) in forward and backward manner through a sliding window of 30s in 1 s steps. To compare trial-by-trial variance, we calculated *V_m_* variance of the trial-by-trial mean in a baseline window (-400 to -300 ms prior to movement) and during a peri-movement window (-50 to +50 ms) for each individual neuron. To calculate the synaptic reversal potential for movement-related synaptic input to L5 PNs, we extracted trial-by-trial baseline *V_m_* and *V_m_* changes (Δ*V_m_*) for the same baseline and peri-movement windows and from the linear fit of Δ*V_m_* vs *V_m_* extracted *E*_*rev*_ at Δ*V_m_*= 0. Membrane potential variance changes in response to thalamic photostimulation were analyzed as mean variance after photostimulation onset (0 to 200 ms after photostimulation onset) normalized to baseline variance (-1 s to 0 before photostimulation onset). For comparison between control trials and trials after Muscimol infusion, we calculated *V_m_* variance from cue aligned trials as the mean variance from cue to median reaction time + 200 ms normalized to baseline variance (-1 s to 0 relative to cue). For the control condition we used all trials prior to thalamic Muscimol injection. For the Muscimol condition we used all trials post Muscimol infusion. Conductance changes were estimated from unfiltered membrane potential recordings using the method described in Berg and Ditlevsen, 2013 (https://de.mathworks.com/matlabcentral/fileexchange/41774-synapticconductance). This method first estimates the total membrane conductance 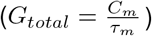 from the membrane time constant extracted from the autocorrelation of the fast *V_m_* fluctuations and the membrane capacitance *C*_*m*_. In a second step, excitatory (*G*_*e*_) and inhibitory (*G*_*i*_) synaptic conductances are then inferred from *G*_*total*_ and the local membrane potential, considering the reversal potential for excitation and inhibition. Because obtaining sufficient behavioral trials under whole-cell recording conditions precluded direct measurement of membrane capacitance and leak conductance, we used *C*_*m*_ = 100 pF and *G*_*m*_ = median (*G*_*total*_)/2 based on previous estimations (Schiemann et al., 2015) and *E*_*leak*_ = -65 mV. Given that we did not correct the membrane potential for the liquid junction potential, we set *E*_*i*_ at - 35 mV and *E*_*e*_ +40 mV relative to spike threshold (-40 mV), which equates to *E*_*i*_= -75 mV and *E*_*e*_ = 0 mV. Importantly, the increase in total conductance is independent of the choice of these parameters as it is directly proportional to *τ*_*m*_. Note that the estimated change in total conductance is independent of these parameter choices, as it is inversely proportional to *τ*_*m*_. Consequently, we report only normalized conductance values throughout the manuscript. To avoid overfitting, we did not adjust parameters for individual neurons if initial estimates yielded non-physiological negative conductances, instead we excluded them from the analysis (4 of 31 neurons).

### Conductance based neuron model

To test how the integration of synaptic conductances affects variability quenching, we developed a conductance-based point neuron model with the following parameters: membrane conductance, *G*_*m*_ = 5 nS; external conductance, *G*_*ext*_ = 5 nS; membrane capacitance, *C*_*m*_ = 100 pF; reversal potential of membrane conductance, *E*_*leak*_ = -65 m; ,reversal potential for inhibitory conductance *E*_*i*_ = -80 mV; reversal potential for excitatory conductance, *E*_*e*_ = 0 mV and variable size of excitatory conductance, *G*_*e*_ and inhibitory conductance, *G*_*i*_. Compound synaptic conductance waveforms for both excitation and inhibition were modeled using two exponential functions with time constant of the rising phase, *τ*_*rise*_ = 0.5s and time constant for the decaying phase *τ*_*decay*_ = 0.4s. Note that these kinetics are not supposed to reflect the individual synaptic kinetics of excitatory and inhibitory postsynaptic currents but represent the time course of the compound synaptic drive during movement initiation. We introduced trial-by-trial variability into the baseline period (preceding the injection of simulated conductance waveforms) by mimicking the effect of different initial E/I ratios on the membrane potential. To do so, we combined the membrane conductance *G*_*m*_ and the external conductance *G*_*ext*_ into a combined conductance *G*_*set*_ (*G*_*set*_ = *G*_*m*_ + *G*_*ext*_) with a reversal potential *E*_*set*_ that we varied from *E*_*set*_ = - 72 mV to *E*_*set*_ = - 58 mV in 1 mV steps. Changing *E*_*set*_ in this way can be thought of as representing different initial E/I ratios in each trial while keeping the overall conductance constant across trials. In this way we were able to test our hypothesis that the magnitude of synaptic conductance input determines the level of variability quenching in a straight forward way without having to account for differences in baseline conductance or differences in fast membrane potential fluctuations from trial to trial. To probe the influence of different magnitudes of synaptic conductances on *V_m_* variability we chose *G*_*e*_ and *G*_*i*_ such that in combination they produced a 2 mV deflection when applied at *E*_*set*_ = -65 mV. To test the effect of input magnitude variability, we varied the maximal variability maxvar from maxvar = 0 to 100% in 10% steps. For each individual trial we randomly chose a trial variability value between -maxvar to +maxvar from a uniform distribution to add to the conductance magnitude. Variability was then calculated from 100 trials at each combination of conductance magnitude and variability size in a time window of 200 ms around the trough of variability under the maxvar = 0 (no variability in synaptic input from trial to trial) conditions. For correlated input variability we used the same random value of trial variability for both *G*_*e*_ and *G*_*i*_, for uncorrelated variability the a trial variability value was drawn independently for *G*_*e*_ and *G*_*i*_.

### Calcium imaging

Raw fluorescence videos were motion corrected using discrete Fourier 2-dimensional-based image alignment (SIMA 1.3.2; Kaifosh et al., 2014). ROIs were drawn manually in Fiji and pixel intensity within each ROI averaged to produce a raw fluorescence time series (F). To remove fluorescence originating from neuropil and neighboring neurons, fluorescence signals were decontaminated and extracted using FISSA (Keemink et al., 2018). Normalized fluorescence was calculated as Δ*F/F* 0 where *F* 0 was calculated as the 5th percentile of the 1 Hz low-pass filtered raw fluorescence signal and Δ*F* = *F − F* 0. All further analyses were performed in custom written scripts in MATLAB. Neurons were considered task-related if they showed significant modulation in a time window of -1 to 1 relative to movement initiation (Kruskal-Wallis test on movement initiation aligned trials, binned in 250 ms windows). We chose this wide analysis window to also include neurons that activate prior to or after movement initiation for an unbiased comparison of activation profiles between different interneuron subtypes. The median onset time of each cell increasing their activity was then calculated by employing a previously published onset detection algorithm using a slope sum function (SSF; Dacre et al., 2021; Zong et al., 2003) with the decision rule and window of the SSF adapted to the calcium imaging data (threshold 10% of peak, SSF window 375 ms, smoothed with a Savitzky Golay filter across 27 frames with order 2 and reported as the median of 10,000 bootstrapped samples to reduce the influence of noisy individual trials). Prior to extracting Δ*F/F* 0 onsets, we verified this algorithm with simulated data thereby accounting for any bias in the onset detection potentially introduced by filtering and/or the decision rule. We then calibrated the onset detection algorithm on the simulated data (100 simulated cells with 50 simulated trials per cell and artificially added noise in each trial matching the noise level in the imaging data) and updated it by a small correction factor.

