## Supplementary Figures S1-S3 for "Thalamic input drives co-timed excitation and inhibition to suppress cortical neuronal variability during movement initiation"

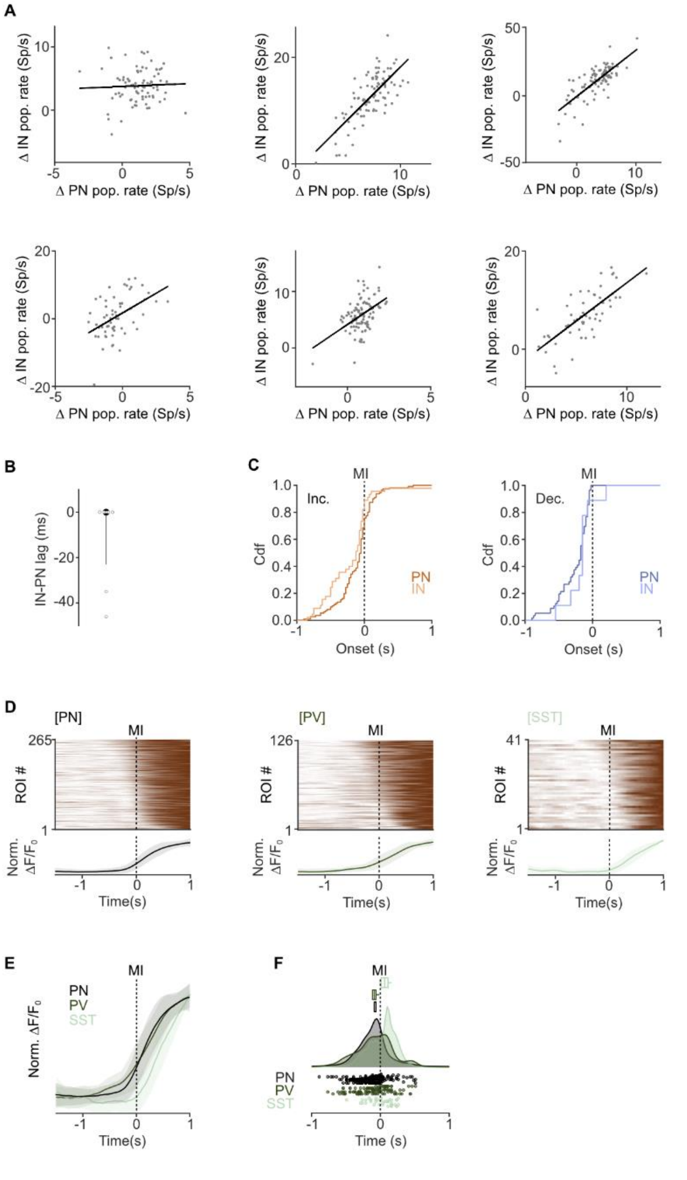

**Figure S1. Timing of pyramidal neuron and interneuron population activity aligned to movement.**
(A) Change in interneuron population firing rate as a function of increasing pyramidal cell population firing rate (N = 6 mice). Grey circles, individual trials; black line, best fit to individual trial data.
(B) Cross-correlation lag between pyramidal neuron and interneuron population activity (N = 6 mice) (related to Figure 1K). Filled circle, median ± 95% confidence intervals; open circles, individual mice.
(C) Cumulative distribution of onset times for pyramidal neuron and interneuron activity changes (*left*, increased; *right*, decreased) relative to movement initiation (MI). Cdf, cumulative distribution function.
(D) *Top*, Normalized across-trial average $\Delta F/F_0$ for pyramidal neurons (PNs, n = 12 sessions, N = 6 mice), pravalbumin- (PV, n = 5 sessions, N = 5 mice) and somatostatin-expressing (SST, n = 6 sessions, N = 4 mice) interneurons aligned to movement initiation (black dashed line). Each row represents the average activity of an individual neuron. *Bottom*, normalized average population $\Delta F/F_0$ ± 95% CI.
(E) Normalized across-trial population average $\Delta F/F_0$ ± 95% CI for pyramidal neurons (PNs), pravalbumin- (PV) and somatostatin-expressing (SST) interneurons aligned to movement initiation (black dashed line).
(F) Distributions of onset times for pyramidal neuron (PN), pravalbumin- (PV) and somatostatin-expressing (SST) interneurons aligned to movement initiation (black dashed line). *Top*, median onset times ± 95% confidence intervals estimated using weighted bootstrap method; probability density across fields of view; median onset times for individual neurons (PN n = 265, PV n = 126, SOM n = 41 neurons).

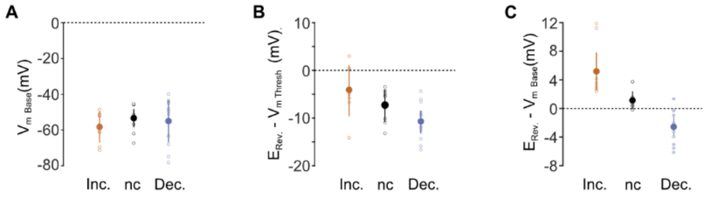


**Figure S2. Subthreshold voltage parameters of layer 5B pyramidal neurons in motor cortex.**
(A) Resting membrane potential ($V_{m\ Base}$) ± 95% CI of pyramidal neurons that increase (red), decrease (blue) or show no change (nc, black) in firing rate during movement (n = 31 neurons, N = 31 mice).
(B) Average distance of the synaptic reversal potential ($E_{Rev.}$) from spike threshold ($V_{m\ Thresh}$) ± 95% CI for pyramidal neurons that increase (red), decrease (blue) or show no change (nc, black) in firing rate during movement. Filled circles, median ± 95% confidence intervals; open circles, individual mice.
(C) Average distance of the synaptic reversal potential ($E_{Rev.}$) from baseline membrane potential ($V_{m\ Base}$) ± 95% CI for pyramidal neurons that increase (red), decrease (blue) or show no change (nc, black) in firing rate during movement. Filled circles, median ± 95% confidence intervals; open circles, individual mice.

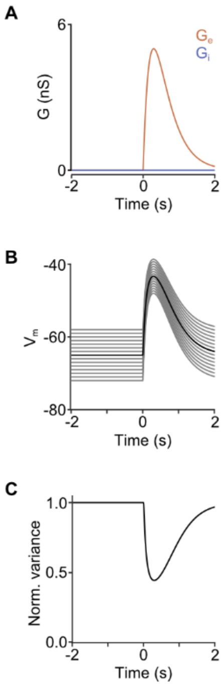


**Figure S3. Membrane potential variance as a function of large changes in excitatory conductance.**
(A) Simulated large excitatory conductance ($G_e$) in the absence of any inhibitory conductance ($G_i$).
(B) Effect of large excitatory conductance across a range of resting membrane potentials ($V_m$).
(C) Normalised variance across trials.
